# Amyloid-ß promotes neurotoxicity by Cdk5-induced p53 stabilization

**DOI:** 10.1101/365437

**Authors:** Rebeca Lapresa, Jesús Agulla, Irene Sánchez-Morán, Juan P. Bolaños, Angeles Almeida

**Author notes:** Corresponding author: Angeles Almeida, Institute of Biomedical Research of Salamanca Calle Zacarías González 2, 37007 Salamanca, Spain.

## Abstract

The p53 tumor suppressor protein, a key regulator of cell apoptosis, has been described to accumulate in affected brain areas from Alzheimer’s disease (AD) patients. However, whether p53 plays any role in AD pathogenesis remains unknown. Here, we found that exposure of neurons to oligomers of the amyloidogenic fragment 25-35 of the Aß peptide (Aβ_25-35_) activated Cdk5, which promoted p53 protein phosphorylation and stabilization. Moreover, Aβ_25-35_-mediated mitochondrial dysfunction and neuronal apoptosis were prevented by both genetic and pharmacological inhibition of either p53 or Cdk5 activities. To confirm this mechanism *in vivo*, Aβ_25-35_ was stereotaxically injected in the cerebral right ventricle of mice, a treatment that caused p53 protein accumulation, dendrite disruption and neuronal death. Furthermore, these effects were prevented in p53 knockout mice or by pharmacologically inhibiting p53. Thus, Aβ_25-35_ triggers Cdk5 activation to induce p53 phosphorylation and stabilization, which leads to neuronal damage. Inhibition of the Cdk5-p53 pathway may therefore represent a novel therapeutic strategy against Aβ-induced neurodegeneration.

## Introduction

Alzheimer’s Disease (AD) is a complex multifactorial neurodegenerative disease (Selkoe and Hardy, 2016) and the most common form of dementia in the elderly (Winblad et al., 2016). The progressive degeneration of neurons in selective brain regions involved in learning and memory, primarily hippocampus and cortex, results in gradual decline of cognitive function (Selkoe and Hardy, 2016). The main neuropathological hallmarks of AD are the accumulation of abnormally folded amyloid-β (Aβ) peptide and truncated and hyperphosphorylated tau proteins in senile plaques and neuronal tangles, which are causally related to the progression of the neurodegenerative process (Scheltens et al., 2016). According to the widely accepted amyloid cascade hypothesis, the accumulation of Aβ in the brain -derived from the proteolytic processing of the amyloid precursor protein- is a primary triggering factor of AD pathogenesis as it initiates a molecular cascade of effects leading to neurodegeneration and the subsequent clinical manifestations of dementia (Karran et al., 2011; Selkoe and Hardy, 2016; Winblad et al., 2016). Aβ oligomers appear to be the most neurotoxic species, triggering signaling pathways that involve synaptic dysfunction, Ca^2+^ homeostasis disruption, oxidative stress, and mitochondrial dysfunction, among others, which culminates in neuronal apoptosis (Green and LaFerla, 2008; Haass and Selkoe, 2007; Jarosz-Griffiths et al., 2016). However, the full underlying mechanism of neuronal apoptosis in response to Aβ is still unknown.

The tumor suppressor protein p53 is a key modulator of cellular responses to genome stress and DNA damage, and p53 activation triggers apoptosis in different cell types, including neurons (Culmsee and Mattson, 2005). Furthermore, numerous evidences demonstrate an increase in p53 level and activity in degenerating neurons, which appears to be a common feature of neurodegenerative diseases (Culmsee and Mattson, 2005; Szybińska and Leśniak, 2017). In particular, enhanced p53 level was detected in brain areas undergoing degeneration of AD patients (Hooper et al., 2007; Kitamura et al., 1997) and transgenic animal models of AD (Ohyagi et al., 2005; Szybińska and Leśniak, 2017). Interestingly, p53 expression increases in parallel with intracellular accumulation of Aβ in some degenerating neurons (Ohyagi et al., 2005). However, it is still a matter of debate whether increased levels of p53 in neurons of AD brains are really implicated in neuronal death or whether they are consequence of the adaptive responses of neurons until apoptosis occurs (Merlo et al., 2014; Sajan et al., 2007; Szybińska and Leśniak, 2017; Zhang et al., 2002). Therefore, the nature of the link between p53 and AD remains controversial.

Here we study the function of p53 in Aβ neurotoxicity and the underlying mechanism, trying to identify possible therapeutic targets for AD. We found that oligomers of the neurotoxic fragment 25-35 of the Aß peptide (Aβ_25-35_) (Pike et al., 1995) promoted Cdk5-induced p53 phosphorylation and functional stabilization, both in primary cultured neurons and *in vivo*, leading to mitochondrial disfunction and neuronal apoptosis. Moreover, we reveal that p53 modulates neuronal susceptibility to Aβ toxicity and determines brain damage, as genetic or pharmacological (PFTα) inhibition of p53 activity confers resistance to Aß-induced dendrite disruption and neurodegeneration. Hence, the Cdk5-p53 signaling pathway may be the link that couples Aβ with neuronal apoptosis, providing novel therapeutic targets for AD therapy.

## Results

### Amyloid-ß induces p53 stabilization leading to neuronal apoptosis *in vitro*

To study the molecular mechanism responsible for Aß oligomer-induced neurotoxicity, primary cortical neurons were first exposed to increasing concentrations of the amyloidogenic fragment 25-35 of the Aß peptide (Aβ_25-35_) for 24 h and neuronal apoptosis was measured by flow cytometry. As shown in Fig. 1A, oligomerized Aβ_25-35_ induced neuronal apoptosis in a concentration-dependent manner, reaching the maximum effect at a concentration of 10 µM, which was selected for further experiments. The time-course study revealed that 10 µM Aβ_25-35_ caused both neuronal apoptosis (Fig. 1B) and mitochondrial membrane potential (∆ψ_m_) depolarization (Fig. 1B). Interestingly, the ∆ψ_m_ loss preceded apoptotic death upon the Aβ_25-35_ exposure, indicating that mitochondrial dysfunction is cause -not consequence- of the neuronal apoptotic process, thus confirming the importance for ∆ψ_m_ maintenance in neuronal survival (Almeida and Bolaños, 2001; Veas-Pérez de Tudela et al., 2015; White and Reynolds, 1996).

**Fig. 1.**
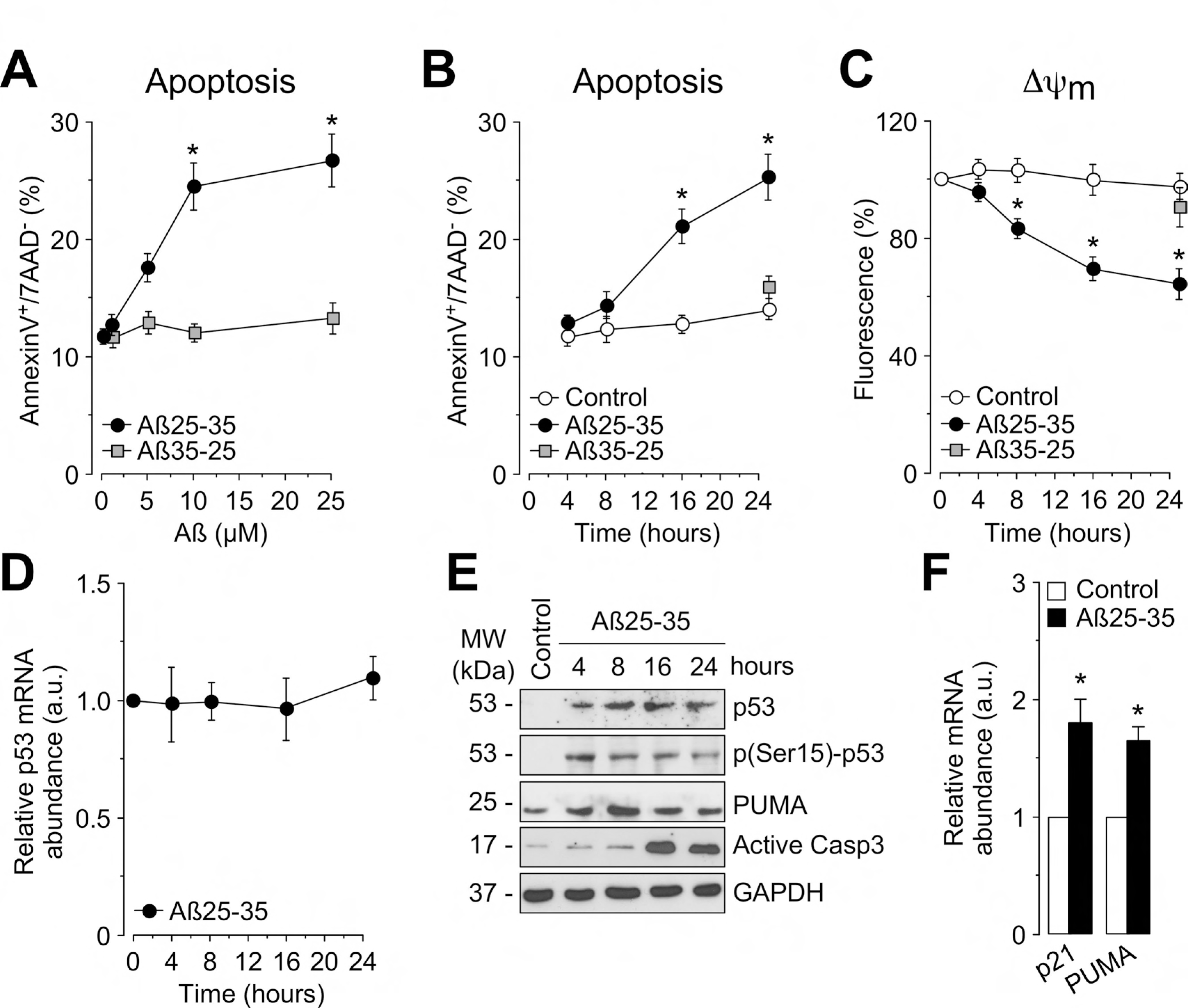
Aß induces p53 stabilization and neuronal apoptosis *in vitro*. (A) Measurement of neuronal apoptosis (annexinV^+^/7AAD^-^ neurons) by flow cytometry reveals that the active fragment Aβ_25-35_ caused a dose-dependent increase in neuronal apoptosis at 24 h of incubation; whereas the inactive Aβ_35-25_ did not affect neuronal survival. (B, C) Aβ_25-35_ (10 µM) exposure results in a time-dependent (B) increase in neuronal apoptosis and (C) mitochondrial depolarization. (D) Time course of p53 mRNA levels measured by RT-qPCR shows that p53 mRNA levels remained unchanged during Aβ_25-35_ exposure. (E) Representative Western blot images showing the time course effect of Aβ_25-35_ on p53 stabilization and phosphorylation (pSer15-p53) and level expression of PUMA and active caspase 3 in neurons. (F) The expression of p53 target genes, p21 and PUMA, were measured by RT-qPCR at 4 h of Aβ_25-35_ exposure. Data are expressed as mean ± SEM from 3-4 different neuronal cultures. *p<0.05 compared to control.

Given the well-known function of p53 in regulating mitochondrial function and neuronal apoptosis (Culmsee and Mattson, 2005; Gomez-Sanchez et al., 2011; Rodríguez et al., 2016), we next examined the contribution of p53 to Aβ_25-35_-induced neurotoxicity. We observed an accumulation of p53 protein in neurons from 4 h of Aβ_25-35_ exposure (Fig. 1E). However, no increase was observed in p53 mRNA levels (Fig. 1D), revealing that Aβ_25-35_-evoked p53 protein accumulation is because of a post-translational event of p53. Under cellular stress conditions, p53 protein levels raises immediately because of its stabilization by phosphorylation (Lee et al., 2007). Accordingly, we found that Aβ_25-35_ promoted p53 phosphorylation at Ser15 residue, which correlated with the timing of p53 protein accumulation (Fig. 1E). Aβ_25-35_-induced accumulation of p53 was functional as revealed by the increase in protein expression of p53 target PUMA, which plays a key role in mitochondrial-mediated apoptosis (Culmsee and Mattson, 2005). Accordingly, Aβ_25-35_ promoted the activation of caspase-3 at 16 h of exposure (Fig. 1E), which correlates with the profile of apoptosis induced by Aβ (Fig. 1B). The functional accumulation of p53 was confirmed by the induction of p53 target genes p21 and PUMA (Fig. 1F).

Next, we analyzed the contribution of p53 accumulation to Aβ_25-35_-induced mitochondrial depolarization and neuronal apoptosis by depleting p53 neuronal levels using both small interfering RNA knockdown (sip53) (Fig. EV1A) and gene knockout of p53 (p53^-/-^).

Following 24 h of Aβ_25-35_ exposure, the lack of p53 accumulation in sip53 (Fig. EV1A) and p53^-/-^ (Fig. 2B) neurons, which prevented the induction of p53 protein targets, p21 and Bax (Fig. 2B), significantly protected against neuronal apoptosis (Fig. 2A and 2C) and caspase-3 activation (Fig. 2B). Moreover, p53 loss prevented Aβ_25-35_-induced mitochondrial depolarization in neurons (Fig. 2D). Finally, the p53-dependent Aβ_25-35_ neurotoxicity was confirmed by pharmacological inhibition of p53 transcriptional activity with PFTα, which decreased Aβ_25-35_-induced neuronal apoptosis in a concentration-dependent manner (Fig. 2E). Furthermore, neuronal protection exerted by PFTα was specific and dependent on p53 activity, as the same effect was observed in p53^+/+^ and p53^-/-^ neurons (Fig. 2F). Altogether, these results demonstrate that Aβ_25-35_ promoted p53 phosphorylation and stabilization, which induced the expression of proapoptotic proteins, PUMA and Bax, leading to mitochondrial dysfunction and neuronal apoptosis.

**Fig. 2.**
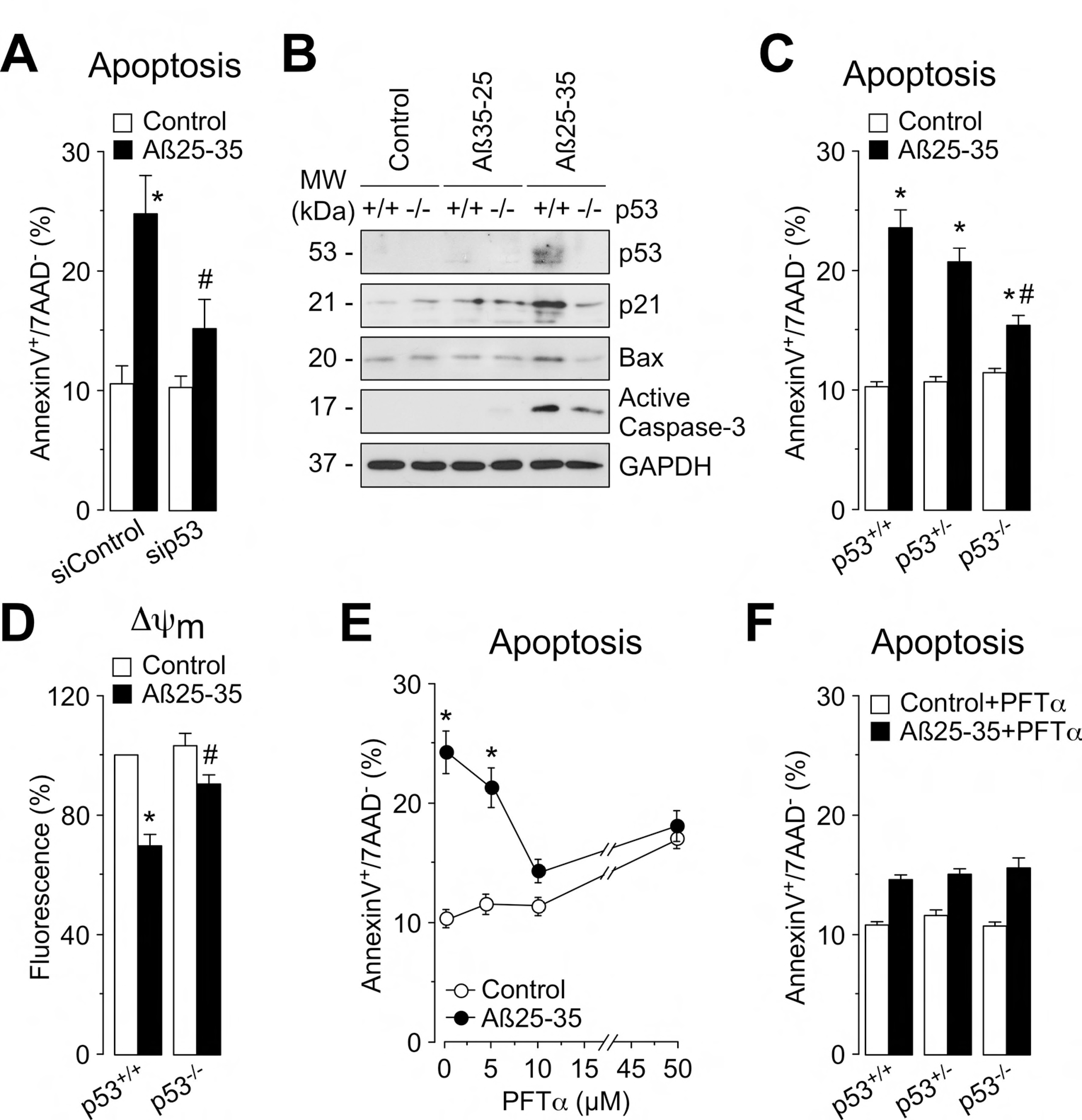
Stabilization of p53 is involved in Aß-induced neuronal apoptosis *in vitro*. (A) Knockdown of p53 by siRNA (sip53) treatment for 2 days significantly decreased Aβ_25-35_ (10µM)-induced neuronal apoptosis (annexinV^+^/7AAD^-^ neurons), in comparison with control (siControl) neurons. (B) Representative Western blot images showing the effect of genetic depletion of p53 (p53^-/-^) on the accumulation of p53 and its targets, p21 and Bax, and the activation of caspase 3, at 24 h of Aβ_25-35_ or Aβ_35-25_ (10 µM) incubation. Flow cytometry analysis shows that genetic deletion of p53 (p53^-/-^) prevented both (C) neuronal apoptosis and (D) mitochondrial depolarization caused by 10 µM Aβ_25-35_ exposure for 24 h. (E) The inhibitor of p53 transactivation activity, PFTα, dose-dependently decreased Aβ_25-35_-induced neuronal apoptosis. (F) The effect of PFTα (10 µM) on Aβ_25-35_-induced neuronal apoptosis was similar in p53^+/+^ and p53^-/-^ mice. Data are expressed as mean ± SEM from 3-4 different neuronal cultures. *p<0.05 compared to control. #p<0.05 compared to Aβ_25-35_-treated siControl (A, D) compared to Aβ_25-35_-treated p53^+/+^ mice.

### Amyloid-ß-induced p53 phosphorylation was mediated by Cdk5 activation

Once demonstrated that Aβ-induced neurotoxicity is dependent on p53 phosphorylation and stabilization, we aimed to identify the kinase involved in the process. It was described that Cdk5, a proline-directed serine/threonine kinase that plays a key role in neuronal degeneration (Maestre et al., 2008; Patrick et al., 1999), phosphorylates p53 on Ser15 under stress conditions, leading to its accumulation (Lee et al., 2007). Cdk5 is persistently activated when bound to its activator p25 (Lee et al., 2000), generated from the Ca^2+^-dependent proteolytic cleavage of p35 (Kusakawa et al., 2000), that accumulates in the brain of AD patients with Alzheimer’s disease (Patrick et al., 1999). As shown in Fig. 3A, Aβ_25-35_ induced the rapid (within 2 h) accumulation of p25. Moreover, Cdk5 activity, as determined by histone H1 phosphorylation, was increased in neuronal extracts exposed to Aβ_25-35_ from 2 h (Fig. 3B), then confirming that Aβ_25-35_ triggered Cdk5 activation in neurons. Next, we measured intracellular free Ca^2+^ concentrations as it promotes calpain activation and consequently p25 formation from p35 (Kusakawa et al., 2000). Aβ_25-35_ induced the rise in Ca^2+^ concentrations in neurons from p53^+/+^, p53^+/-^ and p53^-/-^ mice (Fig. 3C). In all cases, this effect was prevented by the N-methyl-d-aspartate (NMDA) receptor inhibitor, MK801, indicating that intracellular Ca^2+^ influx was mediated by NMDA receptor stimulation. Moreover, Aβ_25-35_ elicited a similar decrease in p35 protein abundance and the accumulation of p25 in both p53^+/+^ and p53^+/-^ neurons (Fig. 3D), indicating that Aβ_25-35_-triggered Cdk5 activation was independent on p53 levels.

**Fig. 3.**
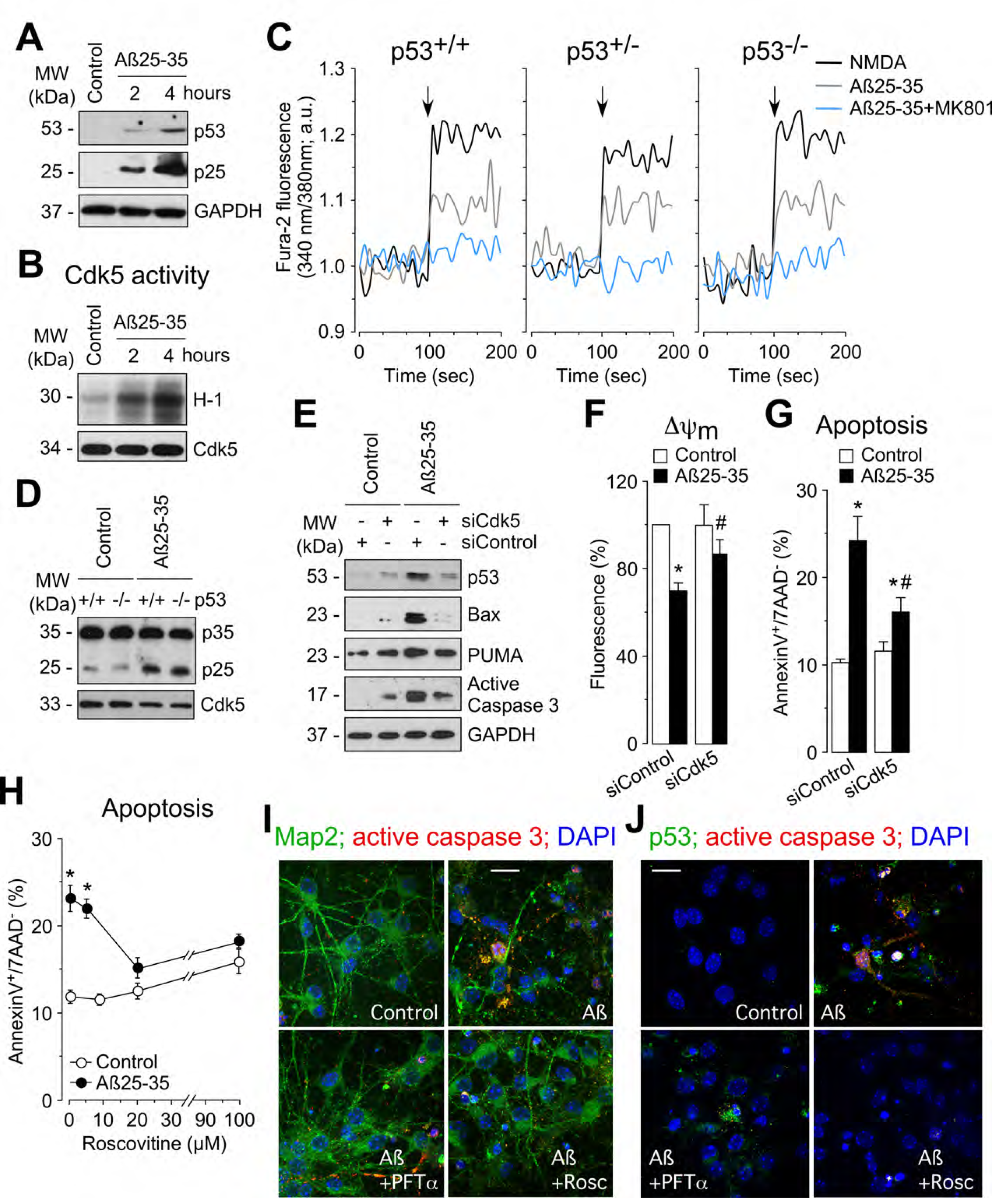
Aß-induced Cdk5 activation triggers p53 accumulation and neuronal apoptosis *in vitro*. Primary cortical neurons were incubated in culture medium in the absence (control) or the presence of oligomerized active Aβ_25-35_ (10 µM), which triggered (A) the accumulation of Cdk5 activator p25 and (B) Cdk5 activation from 2 h of incubation. (C) Neurons from p53^+/+^, p53^+/-^, and p53^-/-^ mice were incubated in culture medium and cytosolic free Ca^2+^ concentrations were measured by fluorimetry before and after NMDA (100 µM) and βA_25-35_ (10 µM) addition. When the experimental condition Aβ_25-35_+MK801 was assayed, the culture medium contained MK801 (10 µM). (D) The formation of p25 from p35 induced by Aβ_25-35_ at 4 h of incubation was not altered by the genetic depletion of p53 (p53^-/-^). (E-G) Neurons were transfected with siRNA against Cdk5 (siCdk5) or luciferase (siControl) for 48 h and then exposed to Aβ_25-35_ (10 µM) for further 24 h. (E) Representative Western blot images reveals that knockdown of Cdk5 (siCdk5) prevented the accumulation of p53 and its targets, Bax and PUMA, and the activation of caspase 3 induced by Aβ_25-35_. Knockdown of Cdk5 (siCdk5) prevented (F) mitochondrial depolarization and (G) neuronal apoptosis induced by Aβ_25-35_. (H) The inhibitor of Cdk5 activity, roscovitine, dose-dependently decreased Aβ_25-35_-induced neuronal apoptosis. (I) Treatment with PFTα (10 µM) and roscovitine (25 µM) prevented the accumulation of active caspase 3 in neurons exposed to Aβ_25-35_ for 24 h, as revealed by immunocytochemistry (Map2, green; active caspase 3, red; DAPI, blue). (J) Aβ_25-35_ induces p53 accumulation in degenerating neurons, as revealed by the co-localization of p53 and active caspase 3 staining, which was prevented by PFTα and roscovitine treatments. Bar: 20 µm. Data are expressed as mean ± SEM from 3 different neuronal cultures. *p<0.05 compared to control; #p<0.05 compared to βA25-35-treated siControl.

Next, we examined whether Cdk5 activation was responsible for the stabilization of p53 upon Aβ_25-35_ exposure. As shown in Fig. 3E, Cdk5 knockdown (siCdk5) (Fig. EV1B) prevented p53 accumulation and the induction of PUMA and Bax proteins caused by Aβ_25-35_, strongly suggesting that Cdk5 increases the stability of the p53 protein. Moreover, knockdown of Cdk5 expression prevented Aβ_25-35_-induced mitochondrial depolarization (Fig. 3F), caspase-3 activation (Fig. 3E), and apoptotic cell death (Fig. 3G) in neurons. The implication of Cdk5 activity on Aβ_25-35_ neurotoxicity was confirmed with the Cdk inhibitor roscovitine, which dose-dependently prevented Aβ_25-35_-caused neuronal apoptosis (Fig. 3H). The effects of Cdk5 knockdown (Fig EV1C and Fig. EV1D) and roscovitine (Fig. EV1E) on mitochondrial function and neuronal apoptosis were similar in p53^+/+^ and p53^-/-^ neurons. Finally, immunocytochemistry images revealed that both roscovitine and PFTα treatments prevented neuronal apoptosis triggered by Aβ_25-35_, as shown by the double-staining for the neuron-specific marker Map2 and active caspase-3 (Fig. 3I). Interestingly, we found that p53 accumulated in degenerating neurons, as shown by the co-localization of p53 and active caspase-3 expression, which was prevented by roscovitine treatment (Fig. 3J). These results support the notion that p53 is phosphorylated by Cdk5 upon Aβ_25-35_ exposure, leading to p53 stabilization and the induction of pro-apoptotic protein expression, Bax and PUMA, which promoted mitochondrial depolarization and neuronal death.

### Amyloid-ß induces p53 stabilization leading to neuronal apoptosis *in vivo*

To demonstrate the physiological relevance of our *in vitro* results, we next aimed to confirm the implication of p53 on Aß neurotoxicity *in vivo*. First, we evaluated the stabilization of p53 in the hippocampal and cortical regions, as they are early targets in AD (Selkoe and Hardy, 2016), of the mouse brain after intracerebroventricular injection of oligomerized Aβ_25-35_. We found that Aβ_25-35_ induced the accumulation of p53 (Fig. 4A) and promoted dendrite disruption, as revealed by the decrease in Map2 staining, in both the hippocampus (Fig. 4B) and cortex (Fig. 4C). Moreover, genetic deletion (p53^-/-^) (Fig. EV1F) or pharmacological inhibition (PFTα) (Fig. 4B and Fig. 4C) of p53 activity significantly improved dendrite disruption induced by Aβ_25-35_.

**Fig. 4.**
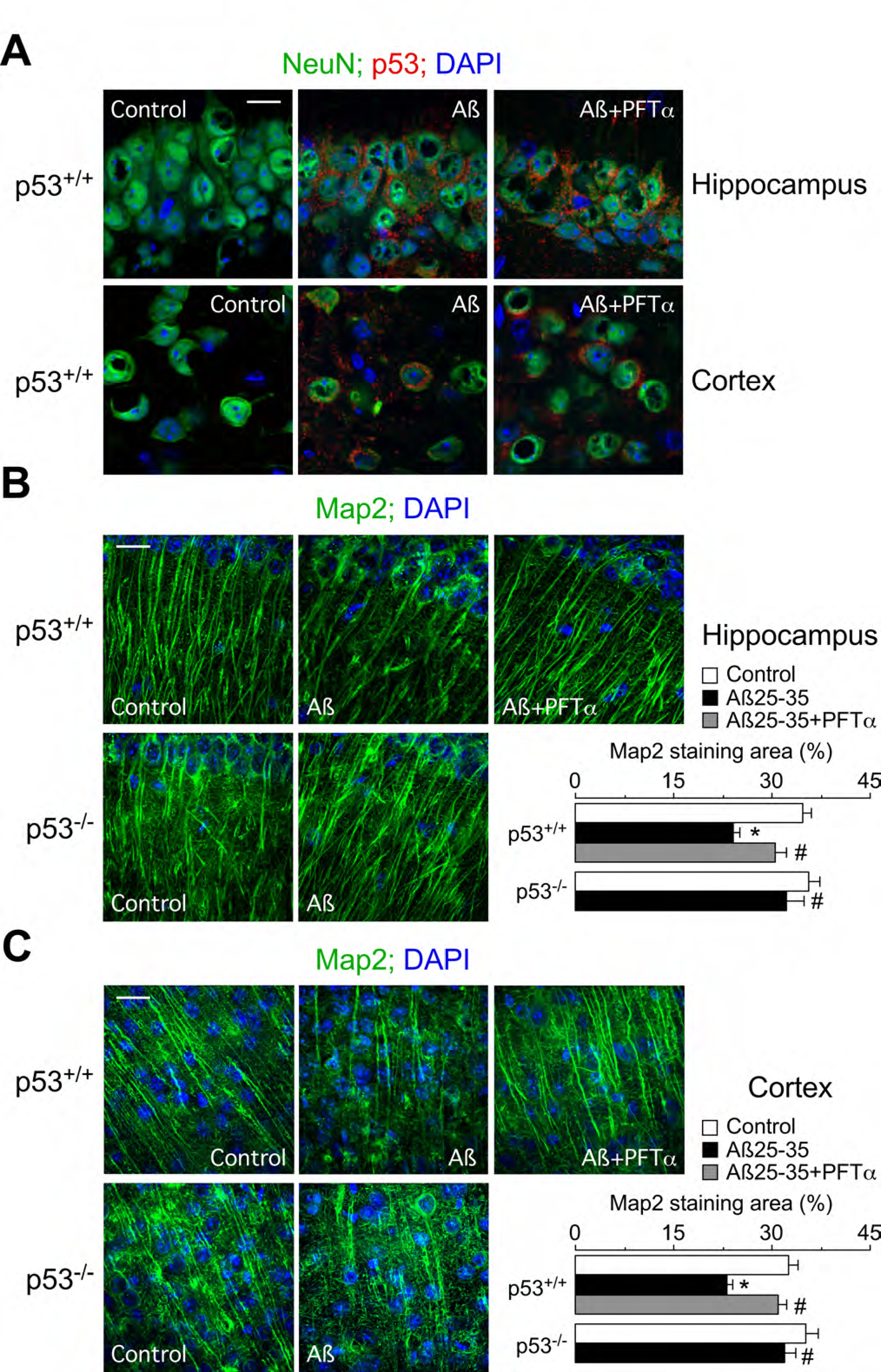
Aß-induced p53 accumulation promotes dendrite disruption *in vivo*. Intracerebroventricular stereotactic injections of 9 nmol Aβ_25-35_ were performed into 12-week-old p53^+/+^ and p53^-/-^ male mice. When indicated, mice were intraperitoneally treated with 20 mg/kg PFTα. (A) Representative images show that Aβ_25-35_ promoted the accumulation of p53 in neurons, as revealed by co-immunostaining for the neuronal marker NeuN and p53 in both the hippocampus (CA1 layer) and cortex (Bar scale: 10 µm; NeuN, green; p53, red; DAPI, blue). (B, C) Representative images show that Aβ_25-35_ promoted dendrite disruption, as revealed by the decrease in Map2 staining in both hippocampus (B) and cortex (C), which was prevented by genetic depletion (p53^-/-^) and pharmacological inhibition (PFTα) of p53 activity (Bar scale: 20 µm; Map2, green; DAPI, blue). Data are expressed as mean ± SEM from 15 different measurements. *p<0.05 compared to p53^+/+^ control; #p<0.05 compared to βA_25-35_-treated p53^+/+^ mice.

Dendrite disruption contributes to the pathology of neurodegenerative disorders, including AD (Cochran et al., 2014). Once demonstrated the implication of p53 on Aβ_25-35_-induced dendrite disruption, we finally studied its possible impact on neurodegeneration. As shown in Fig. 5, double staining of NeuN with TUNEL revealed that Aβ_25-35_-triggered neuronal apoptosis, which was prevented by p53 loss (p53^-/-^) and inhibition of p53 transactivation activity with PFTα treatment. These *in vivo* results are consistent with our findings from cultured neurons *in vitro* and then confirm the relevance of p53 stabilization on Aβ_25-35_-induced neurodegeneration.

**Fig. 5.**
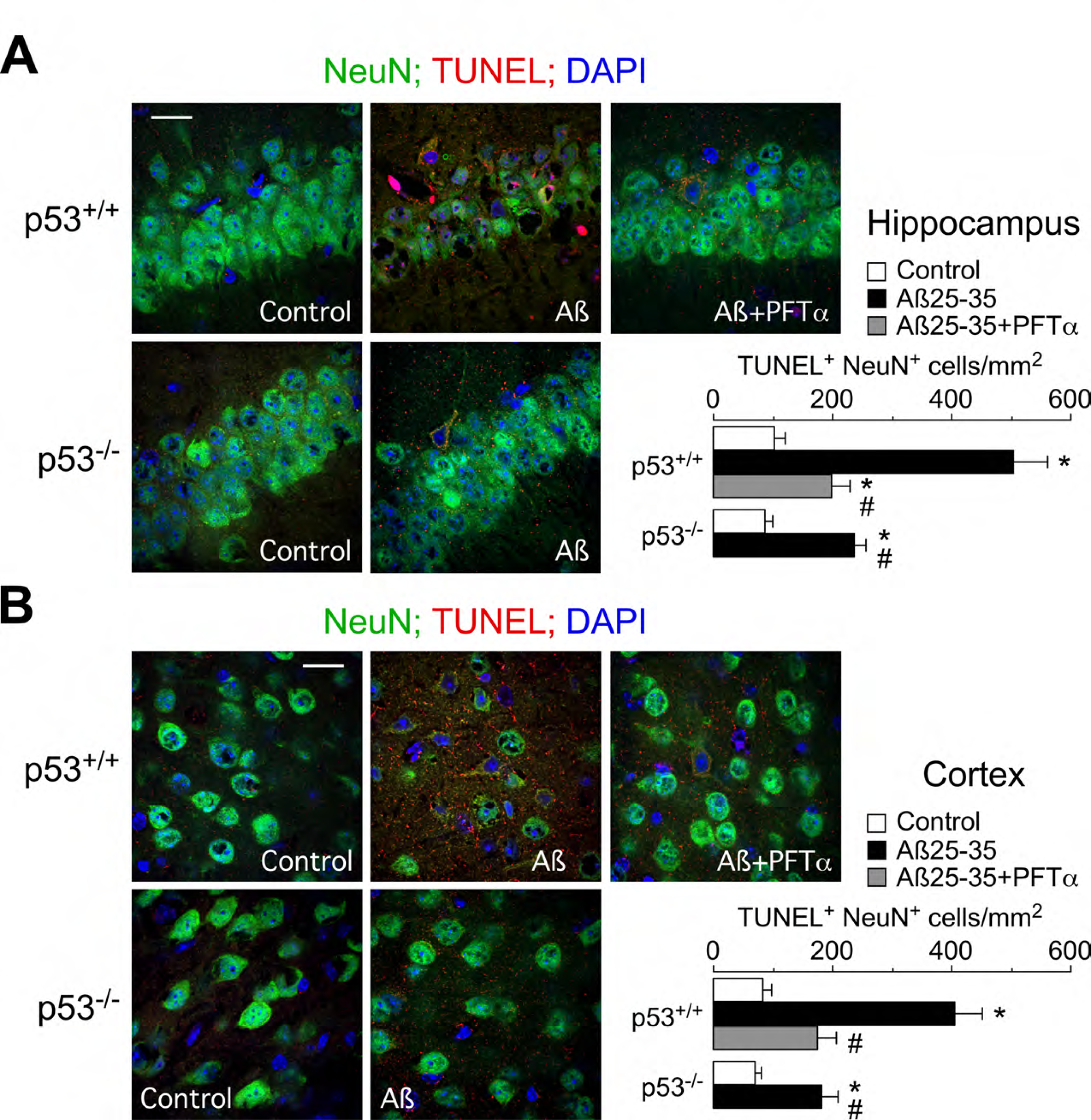
Genetic and pharmacological inhibition of p53 activity prevented Aß-induced neurodegeneration *in vivo*. Intracerebroventricular stereotactic injections of 9 nmol Aβ_25-35_ were performed into 12-week-old p53^+/+^ and p53^-/-^ male mice. When indicated, mice were intraperitoneally treated with 20 mg/kg PFTα. (A, B) Representative images showing that Aβ_25-35_ promoted neuronal apoptosis, as revealed by the double staining of NeuN with TUNEL, in both (A) hippocampus and (B) cortex. Quantification of TUNEL^+^ and NeuN^+^ cells reveals that genetic depletion (p53^-/-^) and pharmacological inhibition (PFTα) of p53 activity prevented Aβ_25-35_-induced neurodegeneration (Bar scale: 20 µm; NeuN, green; TUNEL, red; DAPI, blue). Data are expressed as mean ± SEM from 15 different measurements. *p<0.05 compared to p53^+/+^ control; #p<0.05 compared to βA_25-35_-treated p53^+/+^ mice.

## Discussion

Here, we describe a novel signaling pathway responsible for Aβ_25-35_-induced Cdk5 activation leading to p53 phosphorylation and stabilization, eventually causing mitochondrial dysfunction and neuronal apoptosis. Furthermore, we show that this mechanism operates *in vivo* as either genetic or pharmacological ablation of p53 activity confers resistance against Aß-induced neuronal dendrite disruption and neurodegeneration. This Cdk5-p53 signaling pathway may thus be a causal link coupling Aβ with neuronal apoptosis.

Mechanistically, it is known that activation of p53 requires a post-translational modification that leads to the stabilization of the protein, which in turn inhibits MDM2-mediated p53 ubiquitination and proteasomal degradation (Lee et al., 2007). In good agreement with this, we found that Aβ treatment caused p53 phosphorylation and stabilization that induced the expression of mitochondrial proapoptotic proteins PUMA and Bax, and may also directly interacts with mitochondria (Gomez-Sanchez et al., 2011; Green and Kroemer, 2009), causing mitochondrial damage and neurodegeneration. Furthermore, was found that p53 was accumulated in the degenerating neurons, and that loss of p53, or p53 inhibition with PFTα, protected these cells against Aß toxicity by preventing the induction of Bax and PUMA, mitochondrial dysfunction and caspase activation. Altogether, these results strongly suggest that p53 accumulation in the brain of AD patients (Hooper et al., 2007; Kitamura et al., 1997) may play a key role on neurodegeneration.

Our data also provides novel insights on the molecular mechanisms of the Cdk5 and p53 interplay. Thus, it is known that Cdk5 binds p53 to induce its phosphorylation, which attenuates MDM2-mediated p53 ubiquitylation, thus inducing nuclear localization and accumulation of the protein (Lee et al., 2007). In addition, p25-induced Cdk5 hyperactivation (Lee et al., 2000; Liu et al., 2015) is known to take place in several neurological disorders including AD, and this has been associated with alterations in neuronal cytoskeleton, synaptic loss and, ultimately, neurodegeneration (Hisanaga and Endo, 2010; Shah and Rossie, 2018). Finally, our previous results showed that NMDA receptor over-stimulation in cortical neurons cause p25-mediated Cdk5 hyperactivation (Maestre et al., 2008). Now, we found that Aβ increases NMDA receptor-mediated intracellular Ca^2+^ levels, leading to p25 formation (by p35 cleaving), which in turn triggers Cdk5 activation. Notably, our data also show that p53 stabilization upon Aß treatment is a consequence of the formation of the p25/Cdk5 complex. Intriguingly, elevated levels of both p25 (Patrick et al., 1999; Tseng et al., 2002) and p53 (Hooper et al., 2007; Kitamura et al., 1997) have been detected in the cerebral areas undergoing degeneration in AD patients, hence suggesting a potential link between Cdk5/p25 and p53 accumulation and AD. Altogether, these results strongly suggest that the mechanism underlying neuronal apoptosis caused by Aß may occur, at least in part, as a result of a Cdk5-mediated p53 stabilization process that leads to the transcriptional activation of mitochondrial-derived proapoptotic proteins.

It has been suggested that the pharmacological prevention of p53-dependent apoptotic cascade activation may represent a novel neuroprotective strategy against sudden injury or chronic disease (Esposito and Cuzzocrea, 2010). In fact, treatment with the p53 inhibitor, PFTα, has been shown to reduce neuronal death after traumatic brain injury (Plesnila et al., 2007) and to improve the functional recovery following cerebral ischemia (Luo et al., 2009). Interestingly, PFTα has been shown to reduce kainate-induce excitotoxic death and to prevent mitochondrial and dendrite damage (Neema et al., 2005). In good agreement with these observations, we show that PFTα treatment prevents dendrite disruption and the subsequent neuronal death that follows the intracerebroventricular injection of Aß in mice. Dendrite degeneration and mitochondrial dysfunction are both early pathological events in AD and correlate with cognitive and memory deficits (Cai and Tammineni, 2017). Indeed, mitochondrial energy generation is essential to supporting synaptic function and dendrite integrity (Cai and Tammineni, 2017). Therefore, our data supports the notion that PFTα treatment is a promising novel therapeutic strategy against AD by protecting mitochondrial function and dendrite integrity.

In summary, here we show that Aß-induced Cdk5 activation triggers p53 phosphorylation and stabilization, leading to mitochondrial depolarization, dendrite disruption and neuronal death. This suggests that p53 is an attractive therapeutic target for AD treatment and that the p53 inhibitor, PFTα, may represent a possible therapeutic tool to delaying neurodegeneration in AD patients.

## Materials and Methods

### Primary neuronal culture

Neuronal cultures were prepared from C57BL/6J and p53 null (p53^-/-^, B6.129S2, The Jackson Laboratories, Sacramento, CA, USA) mouse embryo (E14.5) cortices. Animals were maintained in specific-pathogen free facilities at the University of Salamanca, in accordance with Spanish legislation (RD53/2013) under license from the Spanish government and the European Union (2010/63/EU). Protocols were approved by the Bioethics Committee of the Institute of Biomedical Research of Salamanca. All efforts were made to minimize the numbers of animals used. Neurons were seeded at 1.8 x 10^5^ cells/cm^2^ in Neurobasal medium (Invitrogen, Madrid, Spain) supplemented with 2% B27 (Invitrogen) and 2 mM glutamine (Invitrogen), and incubated at 37°C in a humidified 5% CO_2_-containing atmosphere. Half of the culture medium was replaced with fresh medium every 3 days. Neurons were used for the experiments on day 9-10 *in vitro* (Delgado-Esteban et al., 2013).

### Neuronal transfections and treatments

To obtain specific protein knockdown, we used the following small interference RNA (siRNAs; only the forward strand shown): Cdk5, 5´-AAGCCGUACCCGAUGUAUC-3´ (corresponding to nucleotides 859-877) (Veas-Pérez De Tudela et al., 2015); p53, 5´-CCACUUGAUGGAGAGUAUU-3´ (corresponding to nucleotides 955-973) (Delgado-Esteban et al., 2013). In all cases, a siRNA against luciferase (5’-CUGACGCGGAAUACUUCGAUU-3’) was used as control siRNA (siControl). Annealed siRNAs were purchased from Dharmacon (Abgene, Thermo Fisher, Epsom, U.K.). Neurons were transfected with siRNA using Lipofectamine RNAiMAX (Invitrogen), following the manufacturer’s instructions, and used after 24 (sip53) and 48 (siCdk5) hours. According to the degree of protein knockdown shown in the Supplementary Fig. EV1, the efficiency of transfection of siRNAs is estimated to be 80-85%.

The active truncated amyloid-β peptide Aβ_25-35_ (BioNova Cientifica S.L., Madrid, Spain), was dissolved in distilled water at a concentration of 1 mg/mL and then incubated at 37 °C for 3 days to induce its oligomerization (Almeida et al., 2005). Neurons were incubated in culture medium containing oligomerized Aβ_25-35_ (1-25 µM), during the time periods indicated in the Figures. In parallel, neurons were incubated in the presence of the scramble non-aggregable peptide Aβ_35-25_ (BioNova Cientifica S.L.). When indicated, neurons were incubated in the presence of the Cdk5 inhibitor, roscovitine (2-100 µM Rosc; Sigma), or the p53 transactivation inhibitor, PFTα (1-50 µM; Sigma) for 24 h.

### Flow cytometric detection of apoptotic cell death and mitochondrial membrane potential (Δψ_m_)

Neurons were carefully detached from the plates using 1 mM EDTA tetrasodium salt in PBS (pH 7.4) at room temperature and neuronal apoptosis and mitochondrial membrane potential (Δψ_m_) was assessed by flow cytometry. Neurons were stained with annexin V-allophycocyanin (APC; Becton Dickinson Biosciences, New Jersey, USA) and 7-aminoactinomycin D (7-AAD; Becton Dickinson Biosciences) in binding buffer (100 mM HEPES, 140 mM NaCl, 2.5 mM CaCl_2_) to quantitatively determine the percentage of apoptotic neurons by flow cytometry. The annexin V-APC-stained neurons that were 7-AAD-negative were considered to be apoptotic. The Δψ_m_ was assessed using the MitoProbe DilC_1_ (1,1′, 3, 3, 3′, 3′ - hexamethylindodicarbo-cyanine iodide) assay kit for flow cytometry (Life Technologies, Eugene, OR, USA), following manufacturer’s instructions. Neurons were incubated with the dye at 37 °C, for 30 min. Δψ_m_ values were expressed as percentages, using carbonyl cyanide 4-(trifluoromethoxy) phenylhydrazone (CCCP; 10 *μ*M) for 15 min to define the 0% Δψ_m_ value. In all cases, triplicates obtained from four different neuronal cultures were analysed on a FACScalibur flow cytometer (15 mW argon ion laser tuned at 488 nm; CellQuest software, Becton Dickinson Biosciences) (Gomez-Sanchez et al., 2011).

### Quantitative reverse transcription-polymerase chain reaction (RT-qPCR) analysis

This was performed in total RNA samples, purified from neurons using a commercially available kit (Sigma), using the Power SYBER Green RNA-to-CT^TM^ 1-Step Kit (Applied Biosystems, Township, USA). Reverse transcription was performed at 48°C for 30 min, and PCR conditions were 10 min at 95°C followed by 40 cycles of 15 s at 95°C and 1 min at 60°C, using the following forward and reverse primers (0.3 µM), respectively (Thermo Scientific): 5’-CAGCAGCGCGTCCACAGAG-3’and 5’-TCCTGATCCAGGCAATCAC-3’ (*p53 Mmu*); 5’-CGAGAACGGTGGAACTTTGAC-3’ and 5’-CAGGGCTCAGGTAGACCTTG-3’ (*p21, Mmu*); 5’-AGCCCAGCAGCACTTAGAGT-3’ and 5’-ACTCCTCCTCCTCCACACGCA-3’ (*PUMA, Mmu*); 5´-TCAGCAATGCCTCCTGCACCA-3´ and 5´-GCATGGACTGTGGTCATGAG-3´ (*GAPDH Mmu*). The mRNA levels of each transcript were normalized to the *GAPDH* mRNA abundance obtained in the same sample. The relative mRNA levels were calculated using the ΔΔC_t_ method and were expressed as the fold change between sample and calibrator.

### Western blot analysis

Cells were lysed in RIPA buffer (2% sodium dodecylsulphate, 2 mM EDTA, 2 mM EGTA and 50 mM Tris pH 7.5) supplemented with phosphatase inhibitors (1mM Na_3_VO_4_ and 50 mM NaF) and protease inhibitors (100 µM phenylmethylsulfonyl fluoride, 50 µg/ml anti-papain, 50 µg/ml pepstatin, 50 µg/ml amastatin, 50 µg/ml leupeptin, 50 µg/ml bestatin and 50 µg/ml soybean trypsin inhibitor), stored on ice for 30 min and boiled for 5 min. Protein concentrations were determined with the BCA (bicinchoninic acid) method, using bovine serum albumin as a standard (BCA Protein Assay kit, Thermo Fisher Scientific). Neuronal extracts were subjected to SDS-polyacrylamide gel (MiniProtean; Bio-Rad). The antibodies used were anti-p53 (1:2000; 2524, Cell Signaling Technology), anti-p(Ser15)-p53 (1:1000; 9286, Cell Signaling Technology); anti-cleaved caspase-3 (1:2000; Asp175, 9661, Cell Signaling), anti-p21 (1:500; 556431, Becton Dickinson Biosciences); anti-p35 (1:500; Sc-820, Santa Cruz Biotechnology); anti-Cdk5 (1:750; C-8 Sc-173, Santa Cruz Biotechnology); anti-Bax (1:500; Sc-493, Santa Cruz Biotechnology, Heidelberg, Germany); anti-PUMA (1:1000; ab54288, Abcam, Cambridge, UK); and anti-GAPDH (1:40000; Ambion, Cambridge, UK) overnight at 4°C. GAPDH was used as loading control. After incubation with horseradish peroxidase-conjugated goat anti-rabbit IgG (Pierce, Thermo Scientific) or goat anti-mouse IgG (Bio-Rad), membranes were incubated with the enhanced chemiluminescence SuperSignal West Dura (Pierce) for 5 min or Immobilon Western Chemiluminiscent HRP Substrate (Merck Millipore; Darmstadt, Germany) for 1 min, before exposure to Kodak XAR-5 film for 1 to 5 min, and the autoradiograms were scanned (Veas-Pérez De Tudela et al., 2015).

### Cdk5 activity assay

Neurons were lysed in ice-cold buffer containing 50 mM Tris (pH 7.5), 150 mM NaCl, 2 mM EDTA, 1% NP-40, supplemented with the phosphatase and protease inhibitors cited above. After clearing debris by centrifugation, extracts (200 µg protein) were incubated with anti-Cdk5 (1µg) for 4 h, at 4°C, followed by the addition of 30 µl of protein A-sepharose (GE Healthcare Life Sciences) for 2 h, at 4°C. Immunoprecipitates were washed four times in lysis buffer and resuspended in kinase buffer (50 mM Hepes pH 7.5, 10 mM MgCl_2_, 1 mM EDTA and 0.1 mM dithiothreitol) containing 20 µM ATP, 2 µCi of [γ-32P]ATP and histone H1 (1 mg/ml; Sigma). Samples were subjected to SDS-polyacrylamide gel (12%) electrophoresis and transferred proteins were visualized by autoradiography or blotted with anti-Cdk5 (Veas-Pérez De Tudela et al., 2015).

### Cytosolic Ca^2+^ determination using Fura-2 fluorescence

Neurons were loaded with the acetoxymethyl ester form of fura-2/AM (2 μM; Molecular Probes, Eugene, OR, USA) at 37°C for 40 min. After loading, neurons were incubated in dye-free standard buffer (140 mM NaCl, 2.5 mM KCl, 15 mM Tris-HCl, 5 mM D-glucose, 1.2 mM Na_2_HPO_4_ 2H_2_O, 1 mM MgSO_4_, 1 mM CaCl_2_; pH 7.4) for 30 min, to allow the conversion of the dye to its Ca^2+^-sensitive form. Standard buffer was replaced by experimental buffer (140 mM NaCl, 2.5 mM KCl, 15 mM Tris-HCl, 5 mM D-glucose, 1 mM Na_2_HPO_4_ 2H_2_O, 1 mM CaCl_2_; pH 7.4) and stimuli were applied directly to the neurons. Intracellular-free Ca^2+^ concentrations were estimated by recording the Fura-2-emitted fluorescence, using a Varioskan Flash (Thermo Fischer, Vantaa, Finland) spectrofluorometer. Neurons were then excited alternatively at 340 and 380 nm, and fluorescence emission was measured at 505 nm. Results are expressed as absolute increase of the ratio of fluorescence above the basal (resting) fluorescence (previously normalized to the resting fluorescence) after the addition of stimuli.

### Immunocytochemistry

Neurons grown on glass coverslips were fixed with 4% (w/v, in PBS) paraformaldehyde for 30 min and immunostained with rabbit anti-cleaved caspase-3 (1:300; Cell Signaling Technology), mouse anti-p53 (1:200; 2524, Cell Signaling Technology) and mouse anti-Map2 (1:500; Sigma). Immunolabeling was detected using IgG-Cy2 (1:500) or IgG-Cy3 (1:500) secondary antibodies (Jackson ImmunoResearch Inc.). Nuclei were stained with 6-diamidino-2-phenylindole (DAPI, Sigma D9542). Coverslips were washed, mounted with SlowFace light antifade reagent (Invitrogen) and examined under an Olympus IX81 Spinning disk confocal microscope (Olympus®, Tokyo, Japan) (Veas-Pérez de Tudela et al., 2015).

### Experimental *in vivo* model of Amyloid-β toxicity by single Aβ_25-35_ intracerebroventricular injection

Stereotaxic injections were performed as previously done (Rodríguez et al., 2016). Twelve-week-old wt (p53^+/+^) and p53-KO (p53^-/-^) male mice were anesthetized by inhalatory induction (4%) and maintained (2.5%) with sevofluorane (Sevorane; Abbot) in a gas mixture of 70% N_2_O, 30% O_2_, using a gas distribution column (Hersill H-3, Madrid, Spain) and a vaporizer (InterMed Penlons Sigma Delta, OX, UK). Mice were placed in a stereotaxic alignment system (Model 1900, David Kopf Instruments, CA, USA) with digital read out (Wizard 550, Anilam, NY, USA) and complemented with a stereomicroscope (Nikon SMZ 645, Tokyo, Japan) and a fiber optic cold light source (Schott KL1500 compact, Mainz, Germany). Injection was performed into the right ventricle at coordinates: 0.22 mm posterior to bregma, 1 mm lateral to midline, and 2.5 mm ventral to dura, using a 5-µl Hamilton syringe (Microliter 65RN, Hamilton, NV, USA) with a 26 S needle (type 2 tip). Either 4 μL of saline or oligomerized Aβ_25-35_ (9 nmol) were injected using a 5-µL Hamilton syringe with a mini-pump (UltraMicroPump III, World Precision Instruments, USA) and a digital controller (Micro4 UMC4; World Precision Instruments, USA), at a rate of 0.8 µl/min during 5 min. The syringe was left in place for 10 min before slowly retracting it to allow for Aβ infusion and to prevent reflux. Wounds were sutured, and animals were allowed to recover from anaesthesia in cages placed on a 37°C thermostatted plate (Plactronic Digital, 25×60, JP Selecta, Barcelona, Spain). When indicated, animals were intraperitoneally treated with the p53 transactivation inhibitor, Pifithrin α (20 mg/kg; PFTα; Sigma), immediately before Aβ injection.

### Immunohistochemistry

Animals were deeply anesthetized by intraperitoneal injection of a mixture (1:4) of xylazine hydrochloride (Rompun; Bayer) and ketamine hydrochloride/chlorbutol (Imalgene; Merial) using 1 mL of the mixture per kg of body weight, and then perfused intraaortically with 0.9% NaCl followed by 5 mL/g body weight of Somogy’s fixative [4% (wt/vol) paraformaldehyde, 0.2% (wt/vol) picric acid in 0.1 M phosphate buffer, pH 7.4]. After perfusion, brains were dissected out sagittally in two parts and postfixed, using Somogy’s fixative, for 2 hours at room temperature. Brain blocks were then rinsed successively for 10 min, 30 min, and 2 h with 0.1 M phosphate buffer solution (PBS; pH 7.4) and sequentially immersed in 10, 20, and 30% (wt/vol) sucrose in PBS until they sank. After cryoprotection, 10, 20 and 40-µm-thick sagittal sections were obtained with a freezing-sliding cryostate (Leica; CM1950 AgProtect). The 10 and 20-µm sections were placed on microscope slides, whereas the 40-µm slices were collected in 0.05% sodium azide (wt/vol) in 0.1 M PBS. Sections were rinsed in 0.1 M PBS three times each for 10 min and then incubated in: i) 1:1000 anti-NeuN (A-60; Merck Millipore), 1:500 anti-Map2 (AP-20; Sigma-Aldrich) or 1:200 anti-p53 (2524, Cell Signaling Technology) in 0.2% Triton X-100 (Sigma-Aldrich) and 5% goat serum (Jackson ImmunoResearch) in 0.1 M PBS for 72 h at 4 °C; ii) fluorophore-conjugated secondary antibodies (Jackson ImmunoResearch) in 0.05% Triton X-100 and 2% goat serum in 0.1 M PBS for 2 h at room temperature; and iii) 0.5 µg/mL DAPI in PBS for 10 min at room temperature (Bobo-Jiménez et al., 2017). After rinsing with PBS, sections were mounted with Fluoromount (Sigma) aqueous mounting medium.

Confocal images were taken with a scanning laser confocal microscope (“Spinning Disk” Roper Scientific Olympus IX81) with three lasers 405, 491 y 561 nm and equipped with 63× PL Apo oil-immersion objective for high resolution imaging and device digital camera (Evolve; Photometrics, Tucson, USA).

The dendrite integrity in the cortex and hippocampus was assayed by analyzing the density of Map2-positive dendrites in three sections per animal. Fluorescence 8-bit images were acquired as z stacks and were exported into ImageJ in tiff format for processing. Images were converted to grayscale 8-bit images and brightness/contrast was adjusted using the ImageJ “auto” function. All Map2-positive dendrites were automatically delineated using the “auto setting threshold” (default method) and “dark background” functions of ImageJ. Thresholded images were subsequently quantified as percent area (area fraction) using the “analyze-measure” function, which represents the percentage of pixels in the image that have been highlighted (% area) (Bobo-Jiménez et al., 2017). Values are mean ± SEM from 15 measurements.

### Terminal deoxynucleotidyl transferase dUTP nick end-labeling (TUNEL) assay

TUNEL assay was performed in 20 µm brain sections, following the manufacturer’s protocol (Roche Diagnostics, Heidelberg, Germany). Brain sections, fixed as above, were preincubated in TUNEL buffer containing 1 mM CoCl_2_, 140 mM sodium cacodylate and 0.3% Triton X-100 in 30 mM Tris buffer, pH 7.2, for 30 min. After incubation at 37 °C with the TUNEL reaction mixture containing terminal deoxynucleotidyl transferase (800 U/ml) and nucleotide mixture (1 μM) for 90 min, sections were rinsed with PBS and counterstained with Cy3-streptavidin (Jackson Immunoresearch Laboratories) (Rodríguez et al., 2016).

### Statistical analysis

Results are expressed as mean ± S.E.M. A one-way ANOVA with a least significant difference *post hoc* test was used to compare values between multiple groups, and a two-tailed, unpaired Student’s t-test was used for two-group comparisons. In all cases, p<0.05 were considered significant. Statistical analyses were performed using SPSS Statistics 24.0 for Macintosh (IBM).

## Acknowledgements

The technical assistances of Lucia Martin, Estefania Prieto, Monica Carabias, Monica Resch and Carmen Castro are acknowledged. The authors are grateful to Bodegas R. López de Heredia Viña Tondonia for funding part of the work. This work was supported by The Instituto de Salud Carlos III (grant numbers PI15/00473; RD16/0019/0018); European Regional Development Fund; Ministerio de Economía y Competitividad (SAF2016-78114-R); European Union’s Horizon 2020 Research and Innovation Programme (grant agreement 686009); and Junta de Castilla y León (grant number IES007P17). RL was funded by Ministerio de Educación, Cultura y Deporte (FPU fellowship AP2010-3655). ISM was funded by Junta Castilla y León and FSE (EDU/346/2013).

## Contributors

Rebeca Lapresa, Irene Sánchez-Morán, and Jesus Agulla performed the *in vitro* experiments and analyzed biochemical data. Rebeca Lapresa and Jesus Agulla performed the *in vivo* experiments. Juan P Bolaños analyzed data and contributed to design the *in vitro* experiments. Angeles Almeida supervised the project, designed the experiments, analyzed data, and wrote the manuscript. All authors contributed to the discussion of the results and read, revised and approved the final version of the manuscript.

## References

Almeida, A., Bolaños, J.P., 2001. A transient inhibition of mitochondrial ATP synthesis by nitric oxide synthase activation triggered apoptosis in primary cortical neurons. J. Neurochem. 77, 676–90.

Almeida, A., Bolaños, J.P., Moreno, S., 2005. Cdh1/Hct1-APC is essential for the survival of postmitotic neurons. J. Neurosci. 25, 8115–8121.

Bobo-Jiménez, V., Delgado-Esteban, M., Angibaud, J., Sánchez-Morán, I., de la Fuente, A., Yajeya, J., Nägerl, U.V., Castillo, J., Bolaños, J.P., Almeida, A., 2017. APC/C ^Cdh1^ -Rock2 pathway controls dendritic integrity and memory. Proc. Natl. Acad. Sci. 114, 4513–4518.

Cai, Q., Tammineni, P., 2017. Mitochondrial Aspects of Synaptic Dysfunction in Alzheimer’s Disease. J. Alzheimers. Dis. 57, 1087–1103.

Cochran, J.N., Hall, A.M., Roberson, E.D., 2014. The dendritic hypothesis for Alzheimer’s disease pathophysiology. Brain Res. Bull. 103, 18–28.

Culmsee, C., Mattson, M.P., 2005. p53 in neuronal apoptosis. Biochem. Biophys. Res. Commun. 331, 761–777.

Delgado-Esteban, M., García-Higuera, I., Maestre, C., Moreno, S., Almeida, A., 2013. APC/C-Cdh1 coordinates neurogenesis and cortical size during development. Nat. Commun. 4, 2879.

Esposito, E., Cuzzocrea, S., 2010. New therapeutic strategy for Parkinson’s and Alzheimer’s disease. Curr. Med. Chem. 17, 2764–74.

Gomez-Sanchez, J.C., Delgado-Esteban, M., Rodriguez-Hernandez, I., Sobrino, T., Perez de la Ossa, N., Reverte, S., Bolaños, J.P., Gonzalez-Sarmiento, R., Castillo, J., Almeida, A., 2011. The human Tp53 Arg72Pro polymorphism explains different functional prognosis in stroke. J. Exp. Med. 208, 429–37.

Green, D.R., Kroemer, G., 2009. Cytoplasmic functions of the tumour suppressor p53. Nature 458, 1127–1130.

Green, K.N., LaFerla, F.M., 2008. Linking calcium to Abeta and Alzheimer’s disease. Neuron 59, 190–194.

Haass, C., Selkoe, D.J., 2007. Soluble protein oligomers in neurodegeneration: lessons from the Alzheimer’s amyloid beta-peptide. Nat. Rev. Mol. Cell Biol. 8, 101–112.

Hisanaga, S., Endo, R., 2010. Regulation and role of cyclin-dependent kinase activity in neuronal survival and death. J Neurochem 115, 1309–1321.

Hooper, C., Meimaridou, E., Tavassoli, M., Melino, G., Lovestone, S., Killick, R., 2007. p53 is upregulated in Alzheimer’s disease and induces tau phosphorylation in HEK293a cells. Neurosci. Lett. 418, 34–37.

Jarosz-Griffiths, H.H., Noble, E., Rushworth, J. V., Hooper, N.M., 2016. Amyloid-β receptors: The good, the bad, and the prion protein. J. Biol. Chem. 291, 3174–3183

Karran, E., Mercken, M., De Strooper, B., 2011. The amyloid cascade hypothesis for Alzheimer’s disease: an appraisal for the development of therapeutics. Nat. Rev. Drug Discov. 10, 698–712.

Kayed, R., Lasagna-Reeves, C.A., 2012. Molecular mechanisms of amyloid oligomers toxicity. Adv. Alzheimer’s Dis. 3, 67–78.

Kitamura, Y., Shimohama, S., Kamoshima, W., Matsuoka, Y., Nomura, Y., Taniguchi, T., 1997. Changes of p53 in the brains of patients with Alzheimer’s disease. Biochem. Biophys. Res. Commun. 232, 418–421.

Kusakawa, G.I., Saito, T., Onuki, R., Ishiguro, K., Kishimoto, T., Hisanaga, S.I., 2000. Calpain-dependent proteolytic cleavage of the p35 cyclin-dependent kinase 5 activator to p25. J. Biol. Chem. 275, 17166–17172.

Lee, J.-H., Kim, H.-S., Lee, S.-J., Kim, K.-T., 2007. Stabilization and activation of p53 induced by Cdk5 contributes to neuronal cell death. J. Cell Sci. 120, 2259–2571.

Lee, M.S., Kwon, Y.T., Li, M., Peng, J., Friedlander, R.M., Tsai, L.H., 2000. Neurotoxicity induces cleavage of p35 to p25 by calpain. Nature 405, 360–364.

Liu, S.-L., Wang, C., Jiang, T., Tan, L., Xing, A., Yu, J.-T., 2015. The Role of Cdk5 in Alzheimer’s Disease. Mol. Neurobiol. 53, 4328–4342

Luo, Y., Kuo, C.C., Shen, H., Chou, J., Greig, N.H., Hoffer, B.J., Wang, Y., 2009. Delayed treatment with a p53 inhibitor enhances recovery in stroke brain. Ann. Neurol. 65, 520–530.

Maestre, C., Delgado-Esteban, M., Gomez-Sanchez, J.C., Bolaños, J.P., Almeida, A., 2008. Cdk5 phosphorylates Cdh1 and modulates cyclin B1 stability in excitotoxicity. EMBO J. 27, 2736–2745.

Merlo, P., Frost, B., Peng, S., Yang, Y.J., Park, P.J., Feany, M., 2014. P53 Prevents Neurodegeneration By Regulating Synaptic Genes. Proc. Natl. Acad. Sci. U. S. A. 111, 18055–18060.

Neema, M., Navarro-Quiroga, I., Chechlacz, M., Gilliams-Francis, K., Liu, J., LaMonica, K., Lin, S.L., Naegele, J.R., 2005. DNA damage and nonhomologous end joining in excitotoxicity: Neuroprotective role of DNA-PKcs in kainic acid-induced seizures. Hippocampus. 15(8):1057–10571.

Ohyagi, Y., Asahara, H., Chui, D.-H., Tsuruta, Y., Sakae, N., Miyoshi, K., Yamada, T., Kikuchi, H., Taniwaki, T., Murai, H., Ikezoe, K., Furuya, H., Kawarabayashi, T., Shoji, M., Checler, F., Iwaki, T., Makifuchi, T., Takeda, K., Kira, J., Tabira, T., 2005. Intracellular Abeta42 activates p53 promoter: a pathway to neurodegeneration in Alzheimer’s disease. FASEB J. 19, 255–257.

Patrick, G.N., Zukerberg, L., Nikolic, M., de la Monte, S., Dikkes, P., Tsai, L.-H., 1999. Conversion of p35 to p25 deregulates Cdk5 activity and promotes neurodegeneration. Nature 402, 615–622.

Pike, C.J., Walencewicz-Wasserman, A.J., Kosmoski, J., Cribbs, D.H., Glabe, C.G., Cotman, C.W., 1995. Structure-activity analyses of beta-amyloid peptides: contributions of the beta 25-35 region to aggregation and neurotoxicity. J. Neurochem. 64, 253–65.

Plesnila, N., von Baumgarten, L., Retiounskaia, M., Engel, D., Ardeshiri, a, Zimmermann, R., Hoffmann, F., Landshamer, S., Wagner, E., Culmsee, C., 2007. Delayed neuronal death after brain trauma involves p53-dependent inhibition of NF-kappaB transcriptional activity. Cell Death Differ. 14, 1529–1541.

Rodríguez, C., Sobrino, T., Agulla, J., Bobo-Jiménez, V., Ramos-Araque, M.E., Duarte, J.J., Gómez-Sánchez, J.C., Bolaños, J.P., Castillo, J., Almeida, A., 2016. Neovascularization and functional recovery after intracerebral hemorrhage is conditioned by the Tp53 Arg72Pro single-nucleotide polymorphism. Cell Death Differ. 24, 1–11.

Sajan, F.D., Martiniuk, F., Marcus, D.L., Frey, W.H., Hite, R., Bordayo, E.Z., Freedman, M.L., 2007. Apoptotic gene expression in Alzheimer’s disease hippocampal tissue. Am. J. Alzheimers. Dis. Other Demen. 22, 319–28.

Scheltens, P., Blennow, K., Breteler, M.M.B., de Strooper, B., Frisoni, G.B., Salloway, S., Van der Flier, W.M., 2016. Alzheimer’s disease. Lancet. 388, 505–517.

Selkoe, D.J., Hardy, J., 2016. The amyloid hypothesis of Alzheimer’s disease at 25 years. EMBO Mol. Med. 8, 595–608.

Shah, K., Rossie, S., 2018. Tale of the Good and the Bad Cdk5: Remodeling of the Actin Cytoskeleton in the Brain. Mol. Neurobiol. 55, 3426–3438

Szybińska, A., Leśniak, W., 2017. P53 Dysfunction in Neurodegenerative Diseases - The Cause or Effect of Pathological Changes? Aging Dis. 8, 506–518.

Tseng, H.C., Zhou, Y., Shen, Y., Tsai, L.H., 2002. A survey of Cdk5 activator p35 and p25 levels in Alzheimer’s disease brains. FEBS Lett. 523, 58–62.

Vaseva, A. V., Moll, U.M., 2009. The mitochondrial p53 pathway. Biochim. Biophys. Acta - Bioenerg. 1787, 414–420.

Veas-Pérez De Tudela, M., Maestre, C., Delgado-Esteban, M., Bolaños, J.P., Almeida, A., 2015. Cdk5-mediated inhibition of APC/C-Cdh1 switches on the cyclin D1-Cdk4-pRb pathway causing aberrant S-phase entry of postmitotic neurons. Sci. Rep. 5, 18180.

White, R.J., Reynolds, I.J., 1996. Mitochondrial depolarization in glutamate-stimulated neurons: an early signal specific to excitotoxin exposure. J. Neurosci. 16, 5688–5697.

Winblad, B., Amouyel, P., Andrieu, S., Ballard, C., Brayne, C., Brodaty, H., Cedazo-Minguez, A., Dubois, B., Edvardsson, D., Feldman, H., Fratiglioni, L., Frisoni, G.B., Gauthier, S., Georges, J., Graff, C., Iqbal, K., Jessen, F., Johansson, G., Jönsson, L., Kivipelto, M., Knapp, M., Mangialasche, F., Melis, R., Nordberg, A., Rikkert, M.O., Qiu, C., Sakmar, T.P., Scheltens, P., Schneider, L.S., Sperling, R., Tjernberg, L.O., Waldemar, G., Wimo, A., Zetterberg, H., 2016. Defeating Alzheimer’s disease and other dementias: A priority for European science and society. Lancet Neurol. 15, 455–532.

Zhang, Y., McLaughlin, R., Goodyer, C., LeBlanc, A., 2002. Selective cytotoxicity of intracellular amyloid ß peptide1-42 through p53 and Bax in cultured primary human neurons. J. Cell Biol. 156, 519–529.

